# MetagenomicsBench: A Verifiable Benchmark for Agentic Metagenomics Analysis

**DOI:** 10.64898/2026.10.05.756819

**Authors:** Zhen Yang, John Connolly, Arjun Banerjee, Shaista Madad

## Abstract

Metagenomics has transformed our ability to characterize microbial communities without relying on cultivation, but translating complex microbiome datasets into reliable biological conclusions remains a major analytical challenge. Although AI agents have improved substantially at software engineering and general data analysis, it remains unclear whether they can reliably analyze and interpret real-world metagenomic data. We introduce MetagenomicsBench, a benchmark of 100 verifiable evaluations derived from published datasets spanning community structure, host-microbiome associations, microbial function, longitudinal dynamics, and microbiome interventions. Each evaluation provides experimental data and a deterministic grader that evaluates recovery of a key analytical or biological result. Across 10,200 trajectories from 34 model-harness configurations, the strongest configuration achieved a 60.0% pass rate. Performance varied across functional domains, data modalities, and biological systems, while greater cost, token usage, and tool use did not consistently correspond to higher accuracy. Trajectory analysis showed that agents often produced technically plausible analyses while making mistakes in problem interpretation, statistical reasoning, and biological interpretation. MetagenomicsBench pro- vides a framework for measuring progress toward AI agents that can reliably make and validate the analytical and biological decisions required for metagenomic research.

## 1 Introduction

Microbes are an essential part of life on Earth. Many of the chemical cycles that sustain life are heavily dependent on and shaped by microbial activity, and microbes play fundamental roles in human health, ecosystem function, and agriculture National Research Council [2007]. Alterations in the human microbiome have been associated with diverse diseases, including type 2 diabetes and pancreatic disorders Sharma and Tripathi [2019], Boicean et al. [2024]. Advances in metagenomics have transformed our ability to study these communities and their genetic diversity without relying on cultivation, providing access to microorganisms that are difficult or impossible to culture. From 16S rRNA sequencing to shotgun metagenomics and long-read sequencing, modern sequencing technologies allow microbial communities to be characterized at unprecedented scale and resolution Woese [1987], Thomas et al. [2012]. However, translating these complex sequencing datasets into reliable biological insights remains a major analytical challenge. As metagenomic datasets continue to grow in size and complexity, robust computational analysis and biological reasoning are becoming increasingly important for understanding microbial diversity, function, and interactions.

LLM-based coding agents have shown strong performance in software engineering tasks, but flawed analysis of microbiome data could cause costly setbacks to research efforts Gihawi et al. [2023]. As the scale and throughput of omics data continue to increase, it is increasingly important to determine whether these agents can help meet the growing demand for biological data analysis, which is rapidly outpacing the number of bioinformaticians qualified to perform it O’Donoghue [2021]. Benchmarks such as scBench and spatialBench have evaluated agents’ performance in the life science domain and their ability to solve short, verifiable tasks Workman et al. [2026, 2025]. Other benchmarks, such as GeneBench Pro and scBench Long, target long-horizon tasks, and assess whether agents can take raw data and perform end-to-end analyses that reflect real-world computational workflows Li and Ho [2026], Diks et al. [2026]. There have also been efforts to assess agent capabilities in the metagenomics space, such as BioSecBench-surveillance, BioSecBench-function, and BioAgent Bench Bhasin et al. [2026], Wang et al. [2026], Fa et al. [2026]. However, the former two focus on pathogen detection, transmission, and function in a biosecurity context, while the latter includes two metagenomics tasks within a broader suite of bioinformatics workflows, providing limited coverage of the field. To our knowledge, there is no comprehensive metagenomics-focused benchmark that evaluates agent capabilities across the diverse challenges of modern metagenomics, including microbial community profiling, microbial ecology, and host-microbe interactions.

Here, we introduce MetagenomicsBench, a verifiable benchmark of 100 evaluations measuring agents’ analysis and interpretation capabilities across metagenomics, using published datasets spanning microbial community profiling, host-associated and environmental microbiomes, and multimodal analyses integrating metagenomics with single-cell and spatial transcriptomics.

Across 9,900 trajectories from 33 model-harness pairs, the top-scoring configurations (Sonnet 5.5 with Claude Code and Pi) achieved pass rates of around 60%, followed by Opus 5 with Pi at 54%. Agents often completed plausible analyses but failed in biological interpretation, including reasoning about taxonomy, functional capacity, confounding, contamination, and assay-specific artifacts. These results highlight that reliable metagenomic analysis requires not only tool use, but also strong biological reasoning and careful interpretation of the data.

## 2 Benchmark Design

All evaluations were constructed using publicly available studies and graded by deterministic graders. Ground truth was derived either by reproducing published results or, in cases where publication claims were not reproducible, by testing multiple plausible methods to form a consensus. Each evaluation underwent guessability and ground-truth robustness checks to ensure that results were robust across alternative analysis paths and to minimize reward hacking. Evaluations then underwent peer review before being included in the benchmark. See Methods for more details.

The 100 evaluations covered five domains: community structure (46), host-microbiome associations (20), microbial function (18), longitudinal dynamics (11), and microbiome interventions (5). These evaluations tested the ability of agents to characterize differences in microbial communities across biological conditions, identify associations between microbial features and host phenotypes, determine microbial functional capabilities and their contributors, analyze changes in microbiomes over time, and assess the effects of interventions such as diet and FMT.

Given the short-horizon design of MetagenomicsBench, we placed greater emphasis on downstream analysis, which provided a rich space for testing biological reasoning in metagenomics. These evaluations spanned diverse scientific questions and failure modes, with representative examples shown in Table 1.

**Table 1:** Representative evaluations across downstream analysis categories.

| Category | Task | Provided Data | Ground Truth |
| --- | --- | --- | --- |
| Taxonomic / Functional Profiling | Characterize the difference in gut antibiotic-resistance reservoirs between pre-FMT recurrent *C. difficile* infection patients and healthy donors, and determine what underlies the difference. | Per-sample ARG detections from shotgun metagenomes, including ARG abundance, antibiotic class, genomic context, contig host assignment, and sample metadata. | Whether the agent distinguishes total resistome abundance from acquired resistance burden and identifies the taxonomic contribution underlying the observed difference. |
| Taxonomic / Functional Profiling | Determine how the gut microbial capacity to synthesize butyrate changes with age. | Shotgun metagenomic pathway abundances, butyrate-synthesis gene abundances, gene-to-pathway annotations, and subject age. | Whether the agent assesses butyrate synthesis across multiple biosynthetic routes rather than inferring functional capacity from a single annotated pathway. |
| Diversity / Differential Analysis | Identify bacterial genera associated with reduced host-cell proliferation in a colorectal carcinoma tissue section. | Spatially matched bacterial genus abundances and host gene-expression profiles from a 10x Genomics Visium tissue section. | Whether the agent identifies convincing genus-proliferation associations while accounting for tissue-context confounding that can produce spurious correlations. |
| Diversity / Differential Analysis | Determine how two *Bifidobacterium* species differ in abundance across delivery mode and antibiotic exposure during the first month of life. | Infant gut shotgun-metagenomic species profiles and sample metadata including age, delivery mode, and antibiotic exposure. | Whether the agent resolves associations at the species level rather than generalizing genus-level patterns across *Bifidobacterium* species. |

## 3 Results

### 3.1 Model and Harness Performance

Across all 100 tasks, the pass rates spanned a 52.3% range (Figure 2A). The lowest scoring model was Nemotron 3.5 Lightning with the Pi harness, scoring 7.7%. The highest scoring model was Claude Sonnet 5.5 with the Claude Code and Pi harness, scoring 60.0%. Sonnet 5.5 with the Claude Code harness scored 10.3% higher than the next non-Anthropic model.

**Figure 1:**
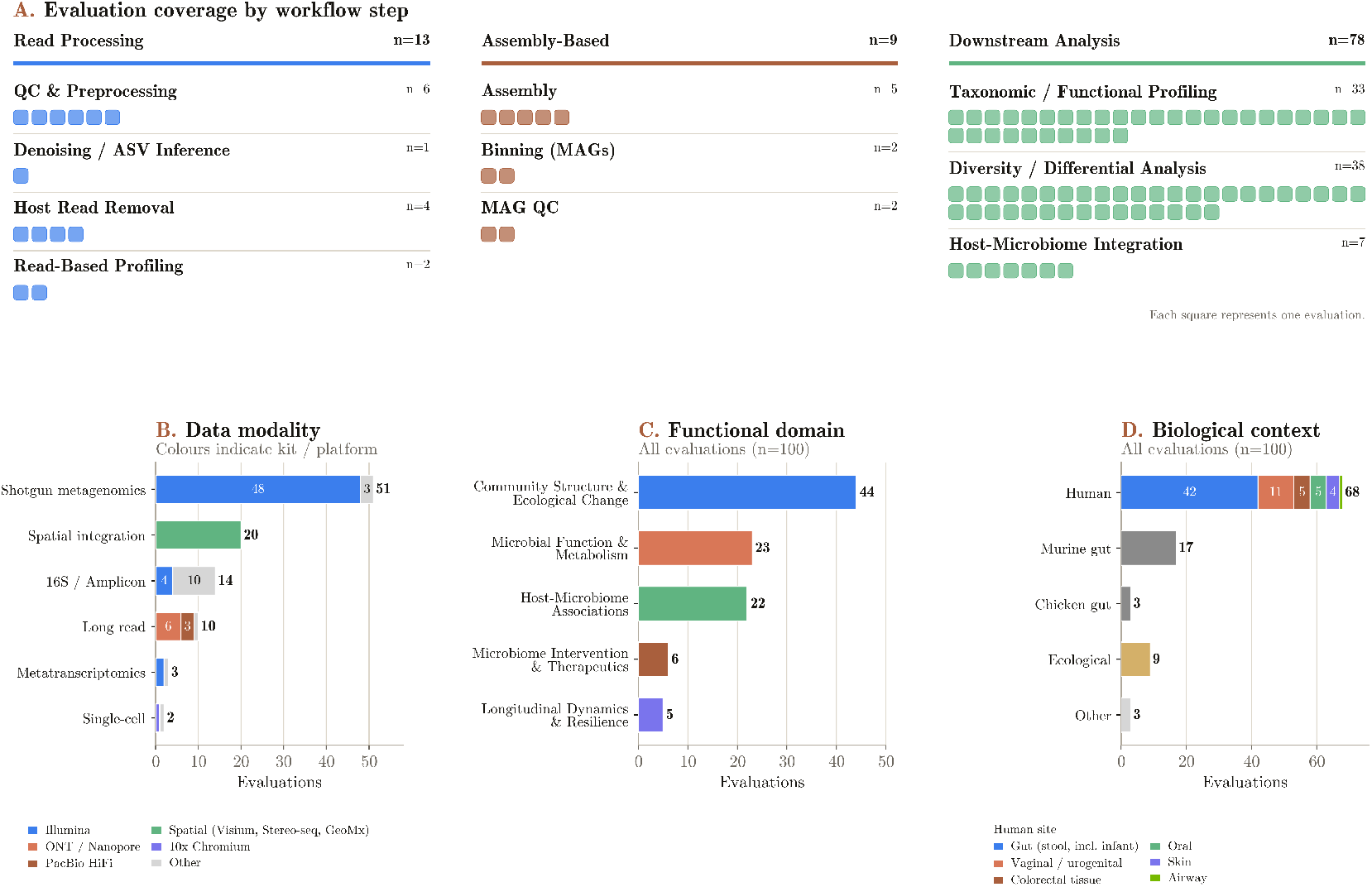
Composition of MetagenomicsBench. Distribution of evaluations across functional domains, sequencing modalities, task taxonomies, biological contexts, and study designs.

**Figure 2:**
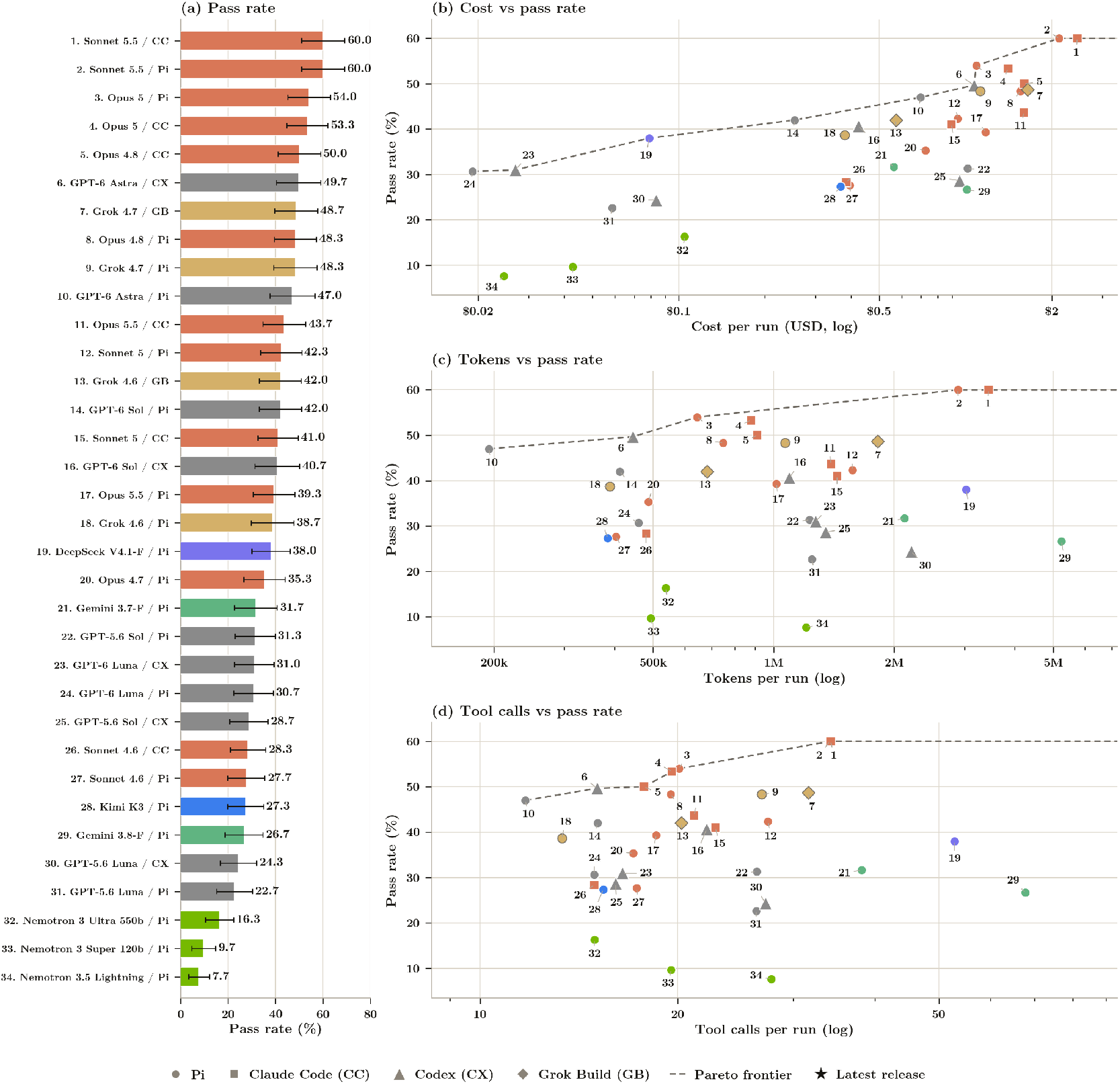
Pass rate and pareto frontiers of MetagenomicsBench. (a) Pass rate of model-harness combinations on . Refusals and timeouts count as failures. Short-horizon rates are the mean over 3 independent trials per task with a 95% eval-clustered Student-t interval; long-horizon rates are the mean over 8 independent trials per task with a 95% eval-clustered Student-t interval. Harnesses are abbreviated GB for Grok Build, Pi for Pi, CC for Claude Code, CX for Codex. A star marks the latest released model from a given provider. (b, c, d) Pareto frontiers of all model-harness combinations in terms of costs, tokens, and turns. Costs are extracted from their respective APIs as of 9/28/26. Colors are based on model family, while shapes are based on harness.

Indeed, the Anthropic models were the highest scoring across the board, with Sonnet-5.5, Opus-5, and Opus-4.8 occupying the 5 model-harness configurations. Conversely, the bottom 3 model-harness combinations were all Nvidia family of models: Nemotron 3 and Nemotron 3.5 Lightning.

Refusals, which were counted as failures, were rare, occupying 2.4% of runs (Appendix A). Refusals were concentrated amongst the Opus 5.5 family of models, which refused 30% of rollouts with the Claude Code harness and 45% with the Pi harness. The next highest refuser were Claude Opus 4.8 with the Claude Code harness and GPT-6-Sol with Codex, refusing 1.33% and 0.67% of trials, respectively.

Timeouts, which were also counted as failures, were rarer, impacting only 0.86% (Appendix A). The model with the most timeouts was Gemini 3.8 Flash with Pi, timing out on 15.33% of runs. The only configurations in the top 5 to have neither refusals nor timeouts were Claude Sonnet 5.5 and Claude Opus 5, both with Pi.

The harness used did not create large differences in pass rate. The median absolute difference between a Pi and other harnesses was 1.0 points for Claude Code, 1.7 for Codex, and 1.8 for Grok Build. There was no overwhelming trend for the direction of the difference. Two of the six Anthropic models scored higher with Pi, as did two of the five OpenAI models.

Harness choice did impact refusal, however. Opus 5.5 scored 43.7% with Claude Code, but 39.3% with Pi. This difference was largely due to refusals, as mentioned above. The two configurations score 62.7% and 71.5%, respectively, on the rollouts that were not refused.

Complementing pass rates, Figure 2B-D plots pass rates against costs, tokens, and tool calls. Mean cost per run varied 128-fold. The cheapest model-harness configuration on the frontier was GPT-6 Luna (less than 2 cents per run), while the most expensive was Claude Code ($2.45 per run). The frontier was jagged; for instance, Opus 5 came within 6 points of the top configuration for less than half the price, while also beating GPT-6 Astra by 4.3% for only 2 more cents.

Much of this difference in cost came from each provider’s token pricing, since the token consumption only varied 27-fold. The heaviest token user, Sonnet 5.5 (3.45M and 2.89M tokens per run for Claude Code and Pi, respectively), also scored the highest. However, Deepseek 4.1 Flash consumed similar amounts of tokens, for a 38.0% pass rate. Below Sonnet 5.5, the link between token spend was less strong. Opus 5 had the third highest score while being in the bottom half of token users, while GPT-6 Astra with Codex was the top quartile of pass rate while being in the bottom quartile of token usage. GPT-6 Astra used the fewest tokens per run while still being in the top half of the pass rate.

The mean number of tool calls per run ranged from 11.7 for GPT-6 Astra with Pi to 67.8 for Gemini 3.8 Flash with Pi, a roughly 5.8x difference. GPT-6 Astra was again the most efficient. With Pi, it made the fewest tool calls of any configuration and needed only 24.9 calls per correct answer, the lowest in the sweep. With Codex, it was the only configuration in both the top quartile of pass rate and the bottom quartile of tool calls. Sonnet 5.5 made 34.2 calls per run to score 60.0%, compared to 19.6 calls for Opus 5 to score 53.3%. Gemini 3.8 Flash made 1.28 times as many calls as the next highest configuration, but scored in the bottom half of the field.

Harness choice did not have a large effect on pass rate, but it had a big effect on consumption. Compared to Pi, the median token usage was 1.21× higher with Claude Code, 1.72× higher with Grok Build, and 2.29× higher with Codex. Grok 4.7 used 1.82M tokens per run with Grok Build and 1.07M with Pi, but the two pass rates were within 0.4 points. GPT-6 Sol used 1.09M tokens with Codex and 0.41M with Pi, and scored 1.3 points lower with Codex.

In general, harness choice did not have a clear impact on tool calls, despite increasing token use. With Claude Code, models made a median of 0.94× as many tool calls as with Pi, but used 1.21× the tokens. Opus 5 showed this most clearly, using 1.37× the tokens and 0.97× the tool calls, and scoring 0.7 points lower. Conversely, Grok Build was the most consistent, raising token usage by 1.75× and 1.70× for its two models.

### 3.2 Performance Across Metagenomics Domains

Model Performances varied across functional domains, data modalities, and biological systems (Figure 3. Within the functional domains, host-microbiome association evaluations generally had lower pass rates across the Opus and GPT frontier model families, while Grok performed best in this category relative to its performance in other functional domains. This advantage was concentrated in a subset of host-microbiome tasks. In matched trajectories where Grok passed and Opus or GPT failed, Grok more often validated key analytical choices before committing to an answer, including accounting for multiple testing, adjusting for potential confounders, and testing the robustness of analysis-specific assumptions. For example, in disease-association tasks, Grok tested the stability of results across prevalence filters rather than relying on a single arbitrary threshold. In spatial tasks, it evaluated whether microbial signals were supported by tissue localization or concordance across multiple probes before interpreting them biologically. In contrast, longitudinal analysis and microbial function were among the categories that consistently achieved higher pass rates across model families.

**Figure 3:**
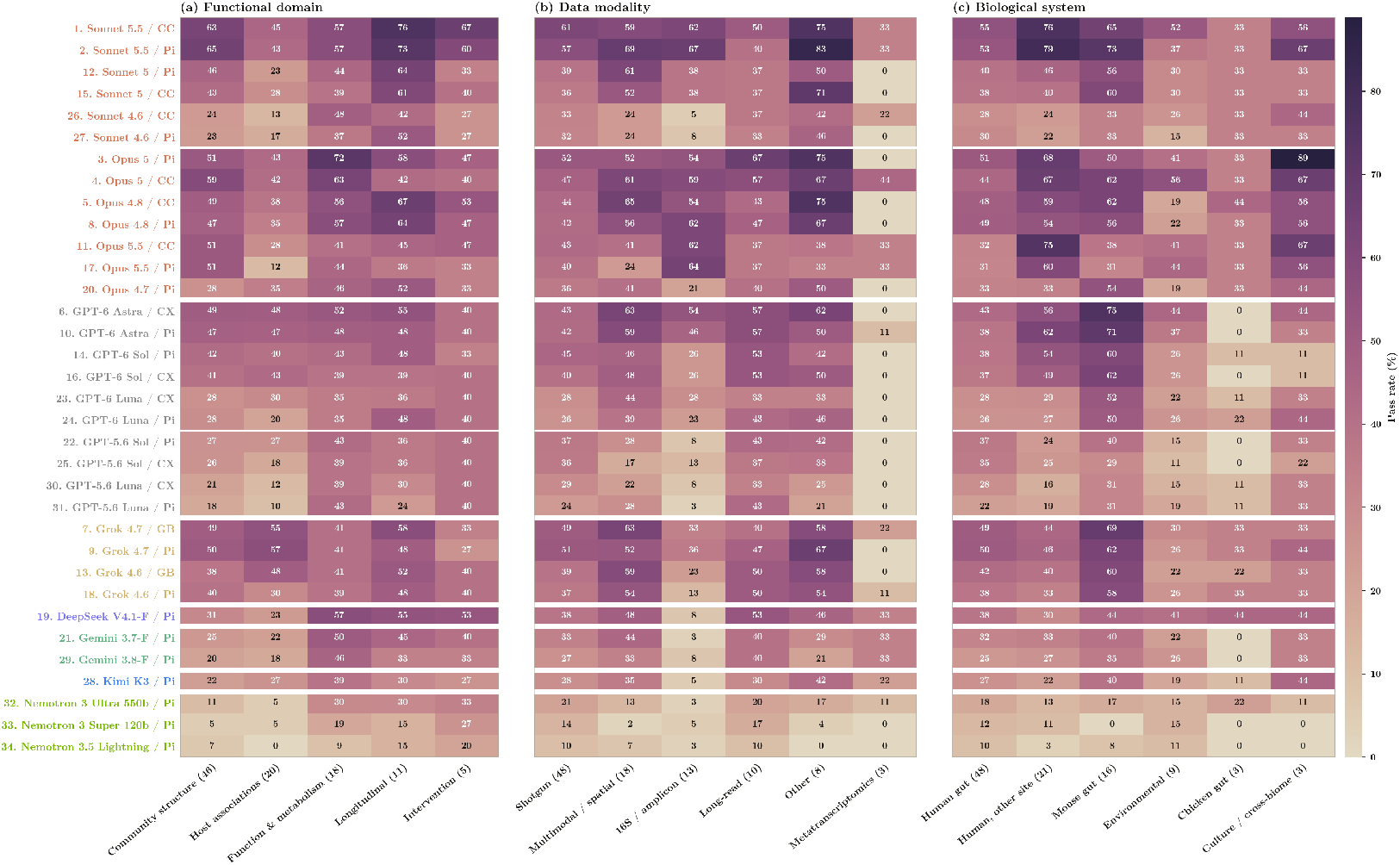
Performance of all model-harness combinations across metagenomics domains, data modalities, and biological systems.

Performance also varied by data modality. Evaluations using 16S data scored highly with the Opus family and Sonnet 5.5, but substantially lower with GPT Sol and Luna. Sonnet 4.6 and GPT 5.6 also scored among the lowest on these evaluations, along with DeepSeek, Gemini, Kimi, and Nemotron. The three evaluations utilizing metatranscriptomics had among the lowest pass rates across nearly all models. Trajectory analysis suggested that these tasks were challenging not simply because of data processing, but because models had difficulty determining what biological conclusions could be supported by transcript-level evidence. Given the small sample size of only three evaluations, however, this pattern may not be representative of model performance on metatranscriptomic analyses as a whole.

Within biological systems, evaluations using the human gut microbiome generally had lower pass rates than those from other human body sites, such as the vaginal, oral, skin, and airway microbiomes, as well as the mouse gut. Failures in human-gut evaluations frequently arose when microbiome-specific analysis had to be integrated with more complex clinical or interventional study designs. Agents often produced analyses that were technically plausible but did not fully account for aspects of the experimental design or the biological interpretation of metagenomic measurements. This pattern suggests that the lower performance may reflect the additional reasoning required to interpret microbiome data in the context of heterogeneous cohorts, longitudinal sampling, and clinical interventions.

Evaluations using the chicken gut microbiome had the lowest average pass rates, although the small number of evaluations limits how representative this result is. Performance on these evaluations also varied substantially across model families. The GPT family had an average pass rate of approximately 7%, compared with 31% for Grok and around 34% for the Opus and Sonnet families.

Together, these results show that performance across metagenomics domains depends not only on the type of data or biological system, but also on how models handle the specific analytical and biological reasoning required by each task.

### 3.3 Failure Modes Analysis

To understand where models were failing across the entire benchmark, Docent’s LLM-as-Judge [Meng et al., 2025] was used to create clusters of failure modes. The traces from all failing runs were normalized, and then passed into Claude Sonnet 5.5, which was instructed to generate an observation, a position where the failure occurred, and evidence. After this, a separate Sonnet 5.5 instance performed adversarial review to verified that the evidence, location, and observations were correct. After this, observations that shared an underlying mechanism into 6 failure modes (Fig 4A). The complete list of mechanisms can be found in Appendix B.

**Figure 4:**
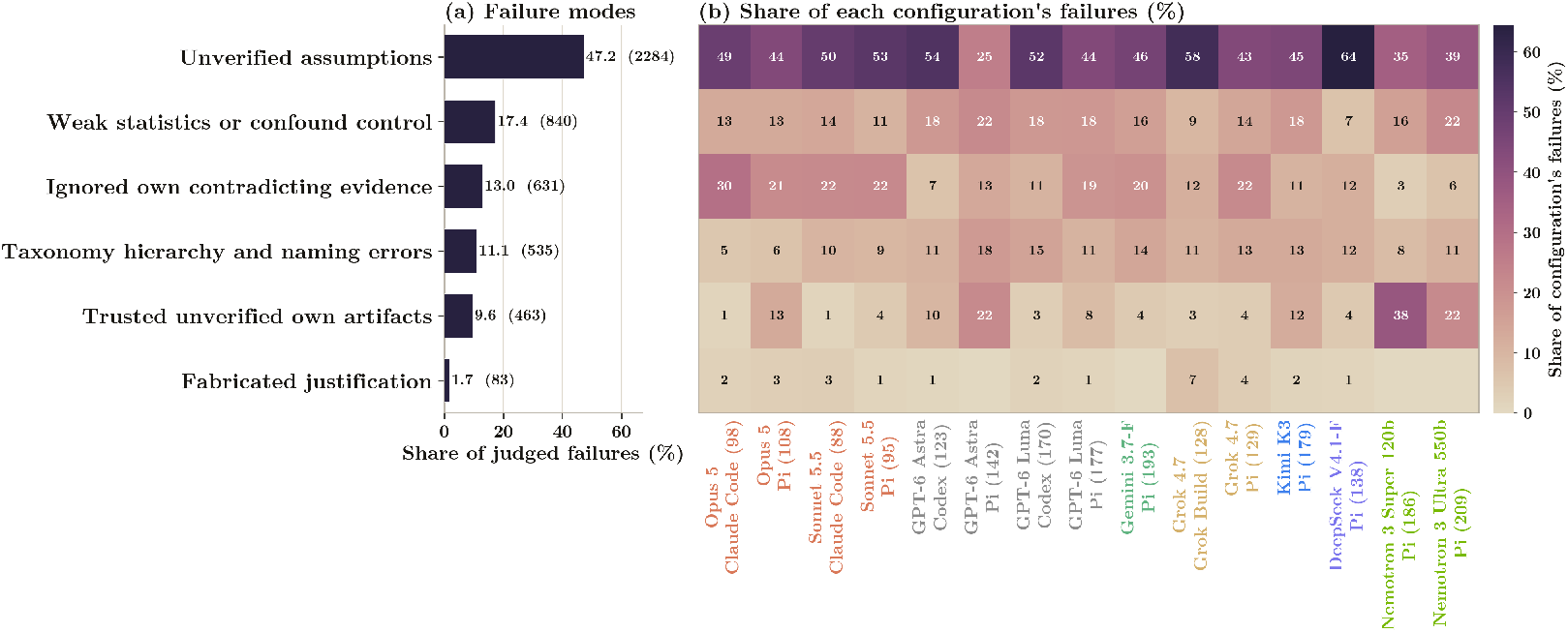
Performance stratified by metagenomic failure modes. The 6 identified failures modes across all failing runs identified by the Docent Judged, ordered by frequency (left). Breakdown of how common identified failrues modes were across a select set of model-harness combinations (right).

The most prevalent failure mode was incorrect problem interpretation: in 47.1% of judged failures the agent encountered what it considered an ambiguous task that a domain scientist would resolve correctly (such as which genes to exclude or a proper detection threshold), and picked otherwise, without recording that a choice had been made. It was the leading mode for every configuration, ranging from 25% (GPT-6 Astra / Pi) to 64% (DeepSeek V4.1 Flash / Pi).

The next most common error was incorrect statistical or confound reasoning (17.4%), which generally took the form of comparing across unequal sequencing depth without normalizing, ignoring a known confound or the non-independence of repeated samples. Next was failure to act on recognized contradictions (13.2%), in which the agent itself surfaced an error in it’s reasoning, but reported the result regardless. Errors in taxonomic hierarchy and nomenclature mechanics followed at 10.9%; these errors included biological reasoning errors like conflating read or row counts with distinct taxon richness. A further 9.7% of failures involved trusting the agent’s own unverified artifacts, such as malformed scripts or validation hardcoded against the target numbers. The final mode, fabricated justification, was rare at 1.8% and described hallucinating justifications for what it believed was a correct answer.

Different model families displayed different failure characteristics (Fig 4B). The Anthropic family of models was significantly more likely to surface problems with model reasoning and ignore it than any other family. 30% of Claude Opus 5 / Claude Code’s failures were ignored contradictions; these detections were quite explicit, noticing batch inflation, noticing confounds, and running deep sensitivity analysis, but reporting results anyways. In general, the Anthropic family had this behavior more than any other model tested. Additionally, Nemotron family trusted it’s own unverified artifacts nearly 2.4x more than the next highest (Kimi-K3 at 12.3%), often answering despite analysis that never ran.

Harness choice also played a factor in failure modes. GPT models run under Codex saw 15.2% more unverified assumptions under Codex than Pi, ranging from a 7 percentage point increase to a 29 point increase. The same comparison held for both Grok Build and Claude Code, increasing unverified assumptions by 9.3 points and 4.4 points on average, respectively. On the other hand, the non-pi harness tended to have less contradiction failures: 7 percent less on Codex and 6.2 percent less on Grok Build. Additionally, Claude Code and Codex had 5.0 and 5.1 percent fewer unverified artifact failures than Pi, respectively.

### 3.4 Computational Resource Awarenes

To see whether agents managed the cost of their own work, the commands in every run were scanned for nine cost-aware acts: checking how many cores and how much memory the machine had, setting a thread count on a tool, timing a command, testing an approach on a subsample before committing to the full data, splitting work into chunks or running it in parallel, launching a job in the background, guarding it with a time limit, killing and restarting it, and inspecting a running process to see whether it was progressing.

In general, cheap forms of cost-awareness were more common than higher-level, complex cost-awareness. Agents checked the machine in 23.7% of runs, but set a thread count in only 9.7% and split or parallelized in only 8.9% of runs (Figure 5a).

**Figure 5:**
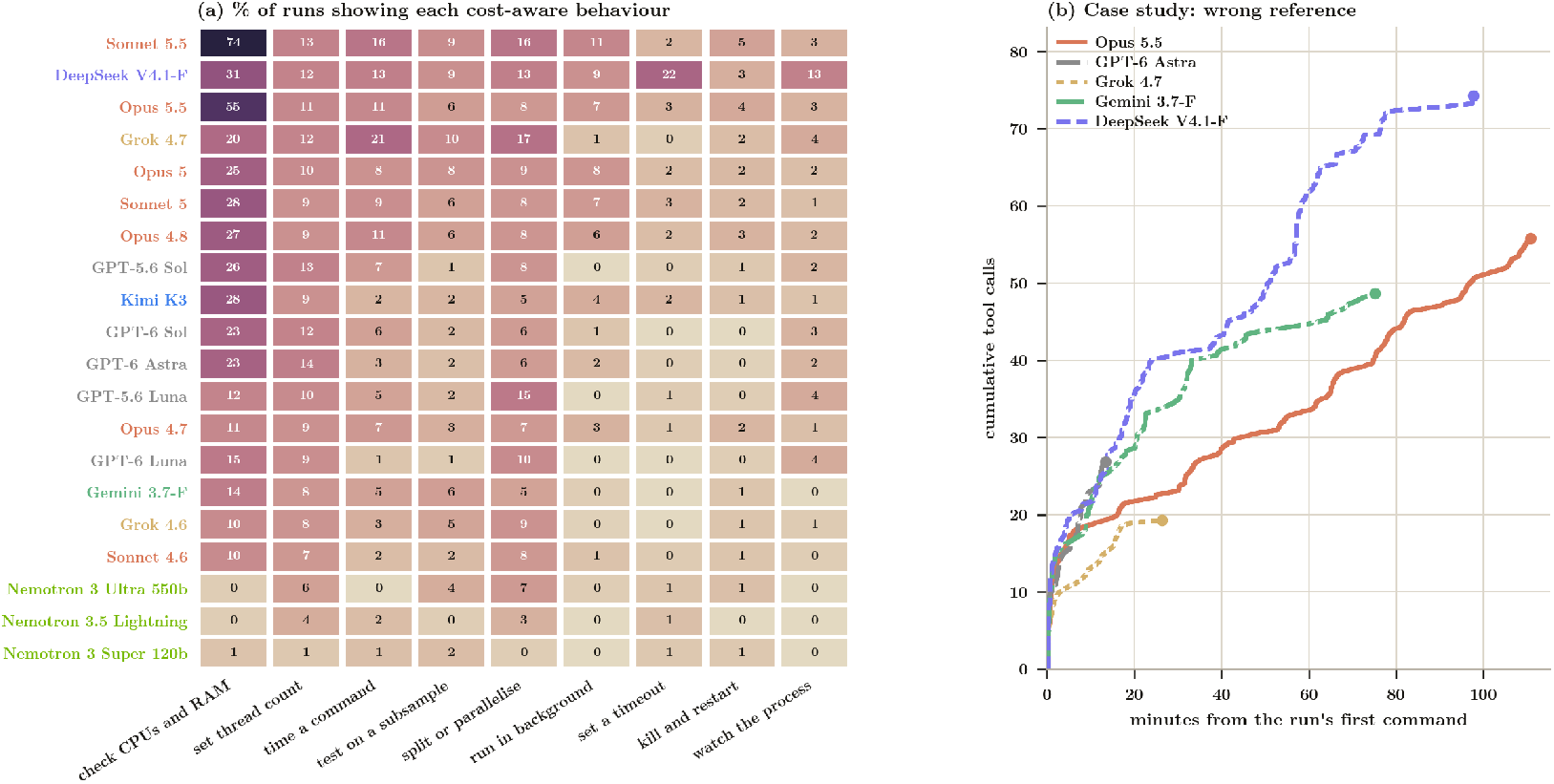
Performance stratified by cost-aware behavior. Breakdown of awarene

There were two distinct outliers, however. Sonnet 5.5 checked the machine in 74% of its runs and Opus 5.5 in 55%; this was in contrast to other model families, which never exceeded 31% for any other configuration. This behavior did not seem to be an Anthropic trait, since Sonnet 4.6 and Opus 4.7 sat at 10% and 11%, but instead a specific behavior in the two newest releases from Anthropic. Moreover, the Deepseek 4.1 Flash was more likely to set timeouts and watch processes than any other model, ensuring that long running commands did not loop forever.

Harness placed a large influence in this behavior as well. For instance, backgrounding appeared in 7% of Claude Code runs and 3% of Pi runs but in none of the Codex or Grok Build runs, so this figure is also a measure of what the harness enabled models to do.

### 3.5 Beyond Endpoint Accuracy: Differences in Host-Reference Selection

Pass rates alone do not capture differences in how agents arrive at a correct answer. We examined a host-fraction quality-control evaluation in which all five representative models reached the ground truth but differed substantially in their analysis trajectories (Figure 5b). The evaluation required agents to determine the appropriate host reference before estimating the fraction of host-derived reads. Purely relying directly on host assignments from taxonomic classification will lead to an incorrect reference choice.

The models differed in how they resolved the reference ambiguity and how much additional validation they performed. Grok 4.7 used a relatively direct comparison between candidate references before applying the selected reference to the full dataset. Gemini 3.7 Flash followed a similarly compact trajectory with less additional validation. Grok 4.7 and GPT-6 Astra reached their final estimates in comparable wall-clock time, although GPT-6 Astra accumulated more tool calls during the analysis. Opus 5.5 performed more extensive validation and required more time, while DeepSeek V4.1-F also reached an accepted answer but performed substantially more exploratory analyses and accumulated the largest number of tool calls.

### 3.6 Adapting to Missing Taxonomic Profiling Tools

Models also differed in how they adapted when standard metagenomics analysis tools were unavailable. We examined an evaluation requiring agents to separately identify the host genus of a plasmid and the most abundant genus in shotgun metagenomic samples. A standard taxonomic profiler was withheld from the sandbox environment, requiring agents to develop an alternative approach for identifying the dominant genus.

Most failed trajectories correctly identified the plasmid host as Prevotella through sequence alignment against the provided plasmid reference database, but failed to identify Actinomyces as the most abundant genus (Figure 6a). This failure was consistent across all three runs of Grok 4.6 and GPT-6 Luna with Pi, while GPT-6 Luna and GPT-6 Sol also failed in two of three runs with Codex (Figure 6b). Several failed agents extracted and classified 16S reads from the shotgun data and used the resulting taxonomic ranking as a proxy for genus-level relative abundance, leading to alternative genera such as Leptotrichia or Centipeda.

**Figure 6:**
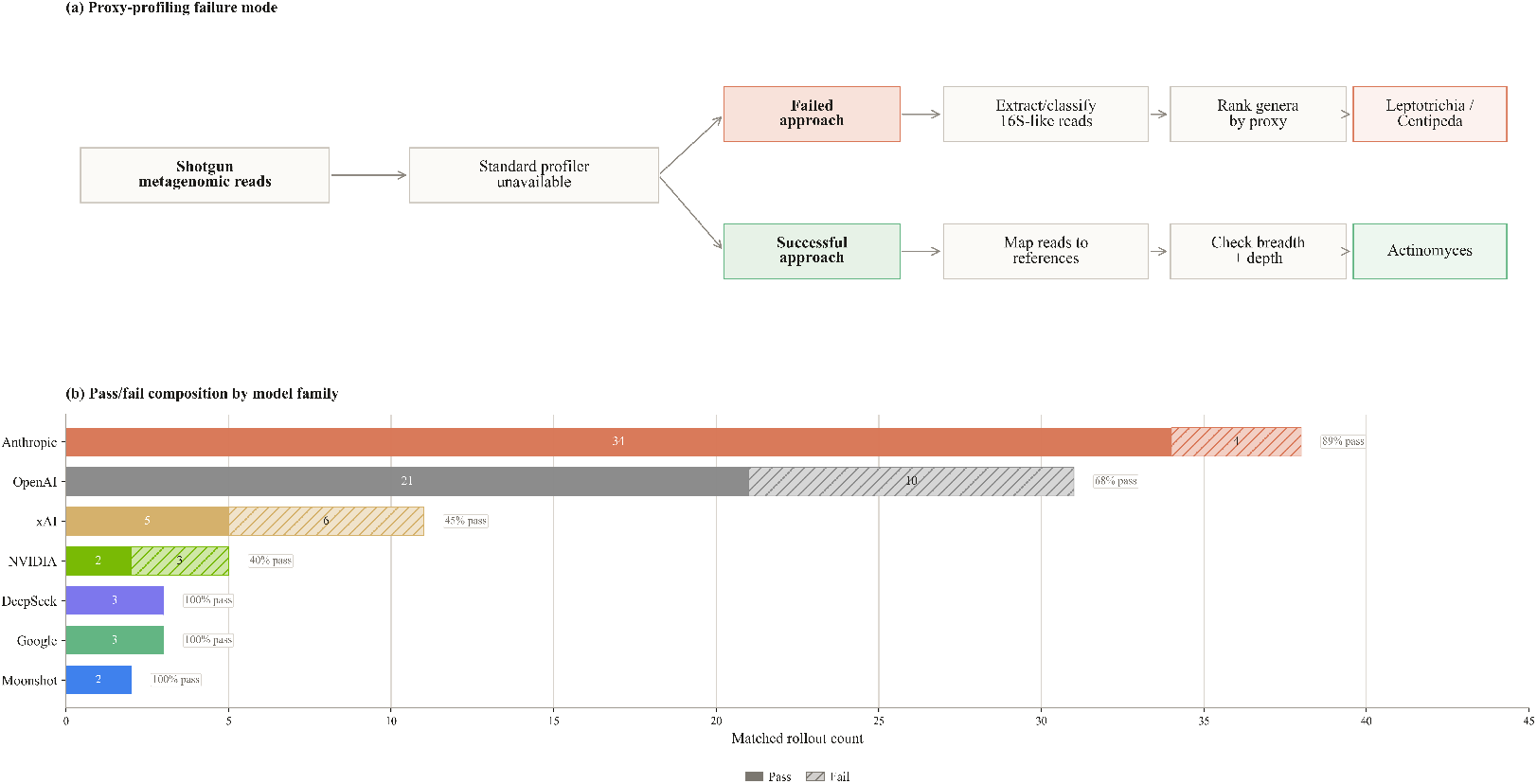
Alternative taxonomic profiling under missing-profiler constraints. **(a)** Divergent analysis strategies used when a standard metagenomic taxonomic profiler was unavailable. Failed trajectories inferred genus-level abundance from 16S reads extracted from the shotgun data, whereas successful trajectories used shotgun read mapping and assessed coverage breadth and depth before identifying *Actinomyces* as the dominant genus. **(b)** Pass/fail composition by model family across 93 model-labeled rollouts for this evaluation.

In contrast, Claude Opus 5, Claude Sonnet 4.6, Claude Sonnet 5.5, GPT-6 Astra, Gemini 3.7 Flash, and DeepSeek V4.1 Flash successfully reached the ground truth. These models treated plasmid host identification and community profiling as separate analyses, identifying the plasmid host by aligning the plasmid sequence against the provided plasmid reference database and estimating the dominant genus from shotgun read mapping, with coverage breadth and depth used to validate the taxonomic signal. For example, Opus 5 found broad coverage across an *Actinomyces oris* reference, providing stronger support for *Actinomyces* as the dominant genus than the rankings obtained from 16S reads alone. These results highlight differences in how models adapt to missing tools and assess the validity of alternative approaches.

## 4 Discussion

We present MetagenomicsBench, a benchmark designed to evaluate whether LLM agents can reliably analyze and interpret metagenomic data across 100 evaluations. The strongest model-harness configuration has 60.1% pass rate, demonstrating that current agents can successfully perform a substantial fraction of these analyses but remain unreliable across the breadth of metagenomic research. Model Performances varied across functional domains, data modalities, and biological systems, with some of the largest challenges arising when metagenomic measurements had to be interpreted in the context of experimental design, clinical variables, or other biological evidence.

Many failures reflected limitations in metagenomic-specific scientific judgment. Incorrect problem interpretation was the most common failure mode, followed by errors involving statistical reasoning and confounding, failure to act on recognized contradictions, and taxonomic reasoning. Agents could therefore produce technically plausible analyses while making analytical or interpretive decisions that changed the biological conclusion. Differences between model families further suggest that similar aggregate performance can arise from different underlying analytical behaviors.

Endpoint accuracy also captures only one aspect of agent performance. Models that reached the same accepted answer could differ substantially in how they selected an analysis, validated intermediate results, and allocated computational effort. The importance of domain-specific reasoning was particularly evident when standard analysis tools were unavailable. When a standard taxonomic profiler was withheld, several models relied on 16S-derived taxonomic rankings that did not reliably represent genus-level abundance, whereas successful models used additional evidence from the shotgun data. Together, these results show that models differ not only in whether they reach the correct answer, but also in how they select and validate analytical approaches. These differences become particularly important when standard tools are unavailable and models must determine whether an alternative method is appropriate for the biological question.

## 5 Limitations

While the benchmark described here represents a genuine test of frontier models’ capabilities, certain aspects of the evaluation structure limited our ability to make holistic conclusions. First, all problems required exact answers (within tolerances). While the strict deterministic grader choice enabled consistent and reproducible scoring, it limited our ability to assess analyses involving context-dependent methodological choices.

This strict grading limitation is particularly relevant to culture-free microbiome analyses, which span extremely diverse ecological contexts and research objectives. As such, the level of standardization is much lower here than in other bioinformatics areas; a 10% microbial prevalence filter may be appropriate when comparing a single cohort of patient stool samples, but may be harder to justify when considering several different groups of soil samples Cao et al. [2021]. In another example, there are several benchmarks in use describing target sequence breadth required to call a population as present in a metagenome, which vary from 1% to 50% Delmont and Eren [2018], Castro et al. [2018], Olm et al. [2021]. Two qualified scientists could very well disagree on what methodological criteria are appropriate for a given situation. We therefore limited our benchmark problems to scenarios with objectively correct or incorrect answers. Future evaluations could instead use rubric-based and partial grading schema to assess how well agents adapt their analyses to the specific context, and justify their methodological choices.

A related limitation concerns the restricted internet access in this benchmark. Internet access was intentionally removed to prevent agents from retrieving answers from the original publications rather than performing the intended analyses. While this may not fully reflect real-world analytical settings, it ensured that performance primarily reflected agents’ analytical capabilities. However, this constraint limited our ability to design questions requiring external resources, databases, or additional tools. Future iterations could use sandboxed environments with a broader, controlled toolset to support more nuanced questions while reducing opportunities for benchmark leakage.

Furthermore, agents were limited to short-horizon tasks on this benchmark, generally consisting of a few analytical steps on a specific data artifact to yield a single set of answers. We did not evaluate models’ abilities to conduct complete, end-to-end metagenomics workflows, as they will likely be asked to do in the real world. As incorrect or sub-optimal decisions compound across a project, long-horizon tasks would likely be more challenging for agents to complete successfully. The pass rates published here could therefore represent an overstatement of models’ true proficiency.

Finally, this initial version of MetagenomicsBench is not an exhaustive survey of culture-free methods and approaches. The evaluations in this dataset principally concern data derived from 16S rRNA amplicon sequencing and shotgun metagenomics, with a small number of tasks involving long read metagenomics, metatranscriptomics, single-cell and spatial transcriptomics. There is limited coverage of metabolomics, metaproteomics, and meta-assembled genomes. MetagenomicsBench primarily focused on bacteria, while other microbial groups, such as viruses, fungi, and their interactions with other microorganisms, remain underrepresented. Host-associated microbial studies included mainly concern the human gut and urogenital microbiota, leaving other important contexts, such as lung microbiomes, skin infections, and other host-associated conditions relatively unexplored. Furthermore, studies investigating microbial metabolism and functional activity at the single-cell level are limited, partly due to the technical challenges associated with capturing microbial DNA and RNA at the single cell level. The areas missing from MetagenomicsBench have received sparse coverage on any published benchmark, and will likely be included in future editions.

## 6 Methods

### 6.1 Benchmark composition and data

The input data of each evaluation were derived from publicly available datasets and then anonymized so agents would not be able to trace the source publication. In cases where only raw FASTQ files were available, they were processed internally, and intermediate files were generated and provided as evaluation inputs if the task itself did not target failure modes in preprocessing steps. Each evaluation was designed around a scientific intent and a set of targeted failure modes. The accompanying notes documented the design principles, targeted failure modes, ground truth justification, and instructions for reproducing the ground truth.

Each evaluation underwent guessability checks by withholding the input data from the agent to make sure the evaluation was not guessable without performing analysis on the data. Other quality checks, such as trajectory analysis and robustness checks of numeric grading tolerances, were also performed to ensure that failure modes were robust and that observed failures reflected the intended scientific failure modes. Evaluations then underwent a final round of peer review, where scientists reviewed each evaluation before acceptance into the benchmark set.

### 6.2 Evaluation Format and Deterministic Grading

Each evaluation was formatted as a JSON file that included a prompt, ground truth, grader, metadata, notes, and the input data required to solve the task. The task prompt specified the answer schema, and the tested agents returned their answers as JSON. The returned answers were then graded against the defined grader. Different types of graders were adopted in MetagenomicsBench, including numeric tolerance, Jaccard similarity, precision and recall, or combinations of these with all of grading, which required agents to get all related fields correct to pass the grader.

### 6.3 Agent Runs and Execution

Each of the 100 evaluations was run three times across model-harness pairs, for a total of 9,900 runs. The harnesses used were pi, claude-code, openai-codex, and grok-build. Models spanned the Claude, GPT, Grok, Gemini, DeepSeek, Kimi, and Nemotron families, with multiple model versions evaluated within each family. Each run was executed in a containerized sandbox with relevant bioinformatics tools installed. The sandbox had no internet access to prevent agents from searching for the source publication or relevant datasets. Each sandbox was provided with 6 CPU cores, 34 GiB of memory, 128 GiB of disk, and a six-hour time limit.

### 6.4 Outcomes and Statistical Analysis

Each run was classified as pass or fail. A run was considered a pass only if all required grading criteria were satisfied. Depending on the grader, partial scores were assigned to provide a more granular failure signal; however, the final outcome was binary. Runs that produced a gradable answer but failed any required criterion were classified as failures. Refusals and runs that reached the six-hour time limit without producing a gradable answer were also counted as failures.

Pass rates were calculated as the number of passing runs divided by the total number of runs for each model-harness configuration. Uncertainty was summarized using 95% Student’s t-intervals computed across per-evaluation mean pass rates, with the evaluation as the sampling unit. This gave each evaluation equal weight in the uncertainty estimate regardless of the number of runs.

## 7 Data Availability

Example evaluations and trajectories will be made available at: https://github.com/latchbio/metagenomicsbench.

## A Run Outcomes

A complete set of run outcomes can be found in Figure 7.

**Figure 7:**
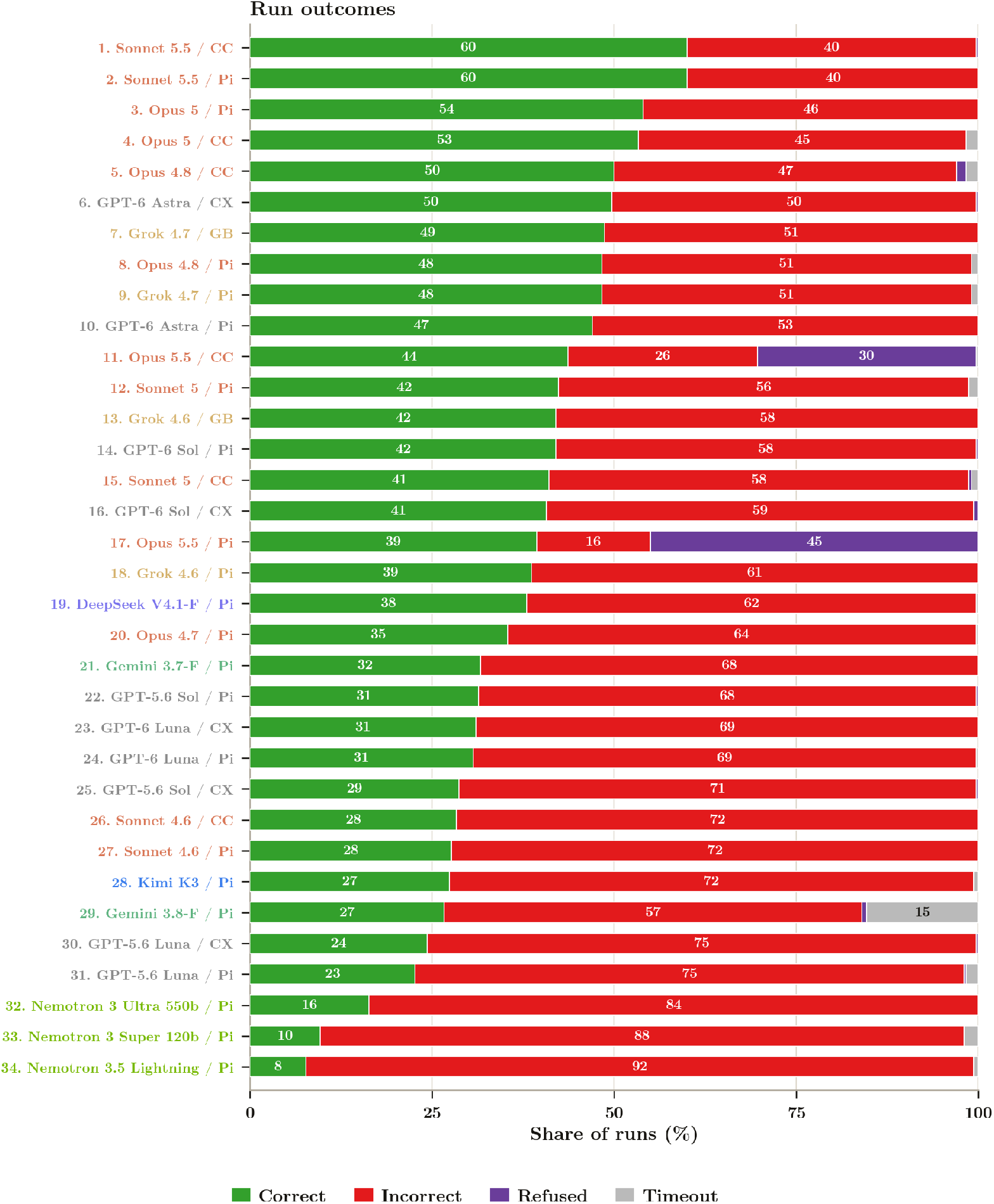
Full breakdown of run outcomes for all model-harness combinations across Metagenomics-Bench, separated into correct, incorrect, timeouts, and refusals.

## B Failure Modes

Full breakdown of failure modes and types identified by the Docent judge pipeline [Meng et al., 2025] (Table 2).

**Table 2:** Failure modes across all judged failing trajectories. Observations from 4,669 failing runs were clustered into 34 mechanisms and grouped into six modes.

| Failure mode | Share | *n* | Description |
| --- | --- | --- | --- |
| Unverified assumptions | 47.1% | 2,200 | The agent met an underspecified term, a threshold, or a metadata field of unstated meaning and picked one reading without recording that a choice had been made. |
| Weak statistics or confound control | 17.4% | 812 | The agent ignored a known confound or non-independence structure, skipped multiple-testing correction, or accepted default tool parameters without validating them. |
| Ignored own contradicting evidence | 13.2% | 616 | The agent surfaced an anomaly, an outlier, or a contradiction in its own reasoning, then reported the result anyway. |
| Taxonomy hierarchy and naming errors | 10.9% | 507 | The agent filtered at the wrong rank, conflated read or row counts with distinct taxon richness, or missed a synonym or reclassification. |
| Trusted unverified own artifacts | 9.7% | 451 | The agent ran a truncated or malformed script, or stopped before any output was validated, and treated the printed numbers as results. |
| Fabricated justification | 1.8% | 83 | The agent hallucinated a rationale for a choice it had already made. |

